# Modular photostable fluorescent DNA blocks dissect the effects of pathogenic mutant kinesin on collective transport

**DOI:** 10.1101/2024.09.06.611758

**Authors:** Tomoki Kita, Ryota Sugie, Yuki Suzuki, Shinsuke Niwa

## Abstract

Intracellular transport is driven by teams of various motor proteins. Advances in DNA nanotechnology have enabled the programmable linkage of different types of motor proteins. In this study, we developed a modular, photostable, fluorescence-labeled tiny DNA origami block (FTOB) for extended observation of collective transport by selected motor combinations. The FTOB is designed as a 4-helix bundle (∼8.4 nm) with densely accumulated fluorescent dyes, minimizing blinking and photobleaching. By designing a pair of connector DNAs, FTOBs are heterodimerized following motor protein attachment using the ALFA-tag/nanobody system. Our system examined the impact of a pathogenic mutant kinesin on its collective movement with wild-type kinesin, clearly observing two distinct behaviors: the team’s velocity was generally governed by the slower mutant but occasionally surged to levels comparable to that of a single wild-type motor. Our photostable, robust, modular FTOB system could serve as a versatile tool for precisely dissecting cooperative cargo transport.

## INTRODUCTION

Intracellular cargo transport is carried out by motor proteins. A variety of motor proteins work together to facilitate efficient and directed transport ^1^. Kinesins, responsible for anterograde transport, typically operate in teams after binding to cargo ^2–4^. These teams can comprise one or multiple types of kinesins, each with distinct mechanical properties, such as velocity, force generation, and directionality ^5–8^. Kinesins also share vesicles with dyneins, the retrograde transporters, suggesting that their tug-of-war mechanism underlies bidirectional transport ^7–11^.

Defects in these transport systems lead to cellular malfunctions. For instance, mutations in neuronal kinesins such as KIF5 (KIF5A, KIF5B, KIF5C) and KIF1A are associated with amyotrophic lateral sclerosis (ALS), hereditary spastic paraplegia, and encephalopathies ^12–18^. These kinesins form homodimers ^19–21^ and are responsible for transporting various cargoes, including membrane organelles, protein complexes, and mRNAs ^17,18^. Most patients with these diseases are heterozygous, possessing both mutated and normal kinesin alleles. This suggests that three distinct types of dimers (i.e., homodimeric wild-type motors, homodimeric mutant motors, and heterodimeric motors composed of one wild-type and one mutant subunit) can bind to the same cargo. While the single-molecule activities of each dimer have been studied ^22–29^, the coordination between different types of dimers on the same cargo remains largely unexplored. To investigate this situation, a methodology that enables accurate control of both the number and types of motor proteins on the cargo is required.

DNA nanotechnology has provided powerful tools that enable the linking of a specific number of different or identical molecular motors via a designed DNA nanostructure ^30–33^. A previous study using such system has shown that combinations of different motor types can lead to significant fluctuations in velocity, likely due to the varying characteristics of the individual motors ^34^. However, frequent blinking and photobleaching of the assembled samples have hindered precise tracking and further analysis of these velocity changes ^34^. One effective solution to this issue is the dense arrangement of multiple fluorescent dyes in a nanometer-scale space, which can be achieved through DNA self-assembly ^35^.

In this study, we developed a connectable, photostable DNA nanostructure, termed the fluorescence-labeled tiny DNA origami block (FTOB). The FTOB is a 4-helix bundle approximately 8.4 nm in size, integrating five or six fluorescent dyes, which minimizes blinking and photobleaching. By designing a pair of connector DNAs, the FTOB can be assembled after attaching the motor protein of interest via the binding between the ALFA-tag and its nanobody ^36^, enabling the formation of a cargo transport complex mimic with a selected combination of motor proteins. Using our modular system, we coupled dimeric KIF5C with disease-associated KIF5C (E237K) and examined their coordinated motion. Our system clearly observed that the rigor mutant primarily impeded the collective movement, yet also allowed short-lived events in which the complex was transiently accelerated to a rate comparable to that of a single wild-type motor.

## RESULTS

### Design and property of FTOB

The FTOB was designed to incorporate six fluorescent dyes, each positioned within a proximity of about 4.5 to 7.1 nm (Fig. 1A). Previous studies have demonstrated that clustering six organic dyes within a few nanometers on a DNA cube assembled from single-stranded tiles (referred to as FluoroCube) significantly increases photon emission and reduces blinking and photobleaching ^35^. Based on these findings, we hypothesized that our FTOB would exhibit similar properties. The FTOB consists of an 84-nt scaffold single-stranded DNA (ssDNA) and four modified 21-nt staple strands. One of these staples contains a functional tag for specific binding to a protein of interest, while the other three are double- or single-fluorescently labeled. To assemble the FTOB after attaching the motor protein, we also prepared a pair of single fluorescently labeled staples, each carrying a 32-nt complementary connector sequence to the other (Fig. 1B). Using a selected set of DNAs, we first prepared six Cy3-labeled FTOB (6xCy3-FTOB) and a pair of five Cy3-labeled FTOBs (referred to as 5xCy3-FTOB-A and 5xCy3-FTOB-a, each containing a complementary connector sequence) (Table. S1 and 2). These FTOBs were self-assembled by thermal annealing and then purified by gel excision.

**Figure 1.**
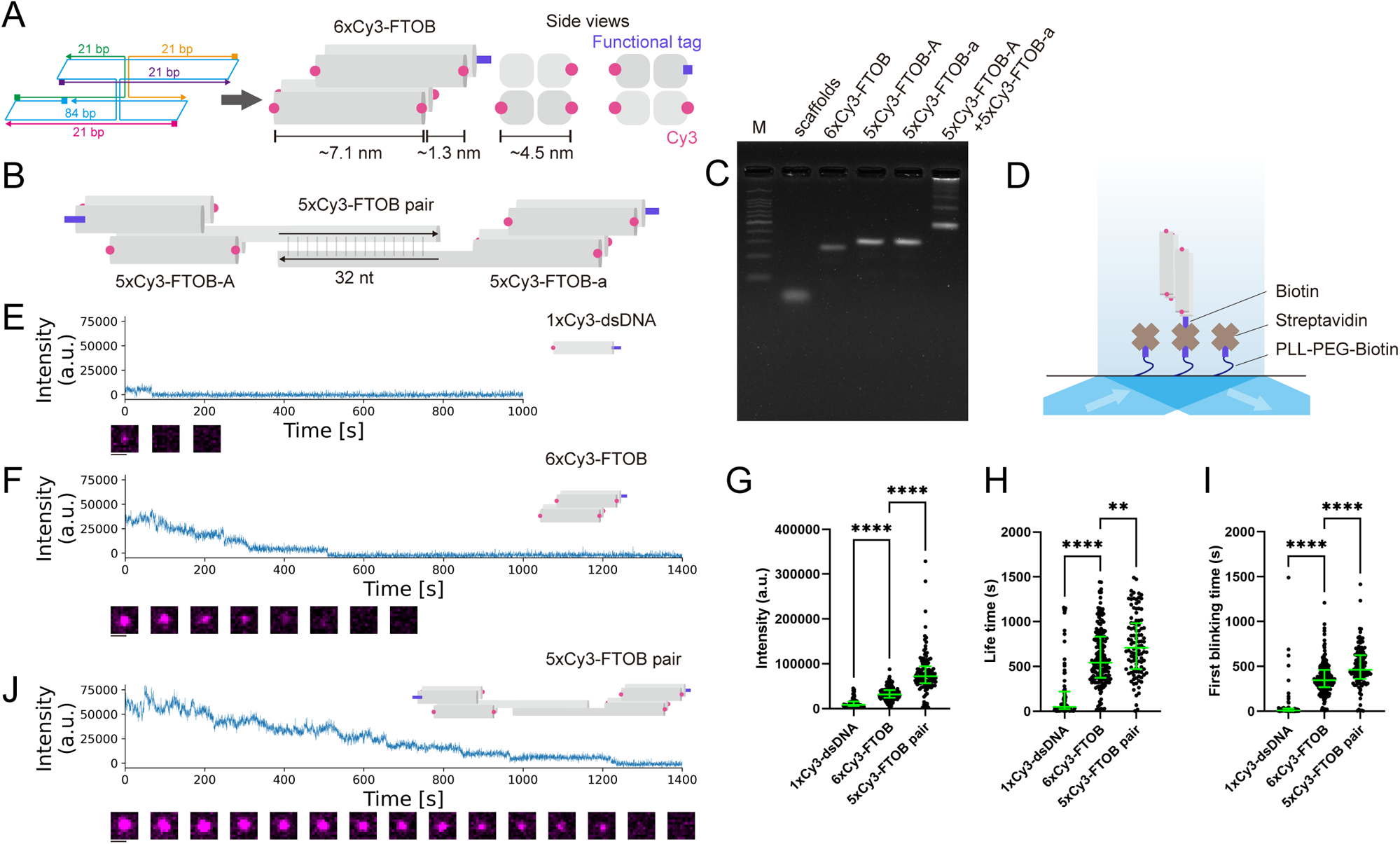
Photophysical properties of FTOBs. (A) Design and schematic illustration of the 6xCy3-FTOB. (B) Schematic illustration of the 5xCy3-FTOB pair. 5xCy3-FTOB-A and 5xCy3-FTOB-a contain 32 nt complementary connector ssDNA and hybridize with each other. (C) Agarose gel electrophoresis analysis of the FTOBs with a biotin-conjugated staple. Lane 1: ladder marker, lane 2: scaffold DNA, lane 3: 6xCy3-FTOB, lane 4: 5xCy3-FTOB-A, lane 5: 5xCy3-FTOB-a, lane 6: 5xCy3-FTOB-A+5xCy3-FTOB-a. (D) Schematic illustration of the experimental setup for measuring photophysical properties of the FTOBs using TIRF microscopy. (E) Example intensity trace of the 1xCy3-dsDNA. Scale bar: 1 μm. (F) Example intensity trace of the 6xCy3-FTOB. Scale bar: 1 μm. (G) Dot plots showing the fluorescence intensity of the 1xCy3-dsDNA, 6xCy3-FTOB, and 5xCy3-FTOB pair. Green bars represent median value and interquartile range. n = 81 (1xCy3-dsDNA), 198 (6xCy3-FTOB), and 122 (5xCy3-FTOB pair). Kruskal–Wallis test followed by Dunn’s multiple-comparison test. ****, p<0.0001. (H) Dot plots showing the lifetime until photobleaching of the 1xCy3-dsDNA, 6xCy3-FTOB, and 5xCy3-FTOB pair. Green bars represent median value and interquartile range. n = 79 (1xCy3-dsDNA), 186 (6xCy3-FTOB), and 105 (5xCy3-FTOB pair). Note that 2, 12, and 17 molecules of the 1xCy3-dsDNA, 6xCy3-FTOB, and 5xCy3-FTOB pair did not undergo photobleaching within 1500 seconds and are not plotted. Kruskal–Wallis test followed by Dunn’s multiple-comparison test. **, p<0.01. ****, p<0.0001. (I) Dot plots showing the first blinking time of the 1xCy3-dsDNA, 6xCy3-FTOB, and 5xCy3-FTOB pair. Green bars represent median value and interquartile range. n = 81 (1xCy3-dsDNA), 198 (6xCy3-FTOB), and 120 (5xCy3-FTOB pair). Note that 2 molecules of the 5xCy3-FTOB pair did not undergo blinking within 1500 seconds and are not plotted. Kruskal–Wallis test followed by Dunn’s multiple-comparison test. ****, p<0.0001. (J) Example intensity trace of the 5xCy3-FTOB pair. Scale bar: 1 μm.

Agarose gel electrophoresis (AGE) of the purified samples confirmed prominent single bands for the series of FTOBs (Fig. 1C). After incubating 5xCy3-FTOB-A and 5xCy3-FTOB-a, successful coupling was demonstrated by the reduction of monomer bands and distinct band shifts in AGE (Fig. 1C).

Next, we examined the fluorescent properties of the FTOB using total internal reflection fluorescence (TIRF) microscopy. The FTOBs with a biotin-conjugated staple were immobilized on a streptavidin-coated glass surface (Fig. 1D). To test whether the 6xCy3-FTOB contained the designed number of dyes, we performed a photobleaching step analysis on the purified samples, confirming that they were predominantly monomers carrying six Cy3 groups (Fig. S1A and B). To quantify brightness and photostability, we compared fluorescent intensity, first blinking time, and lifetime until photobleaching with those of a single Cy3 dye attached to dsDNA (1xCy3-dsDNA). The fluorescent intensity of the 6xCy3-FTOB (median = 3.2 x 10^4^ a.u.) was approximately four times higher than that of the 1xCy3-dsDNA (median = 0.8 x 10^4^ a.u.) (Fig. 1E-G). Additionally, the lifetime (median = 564 s) and first blinking time (median = 345 s) of the 6xCy3-FTOB were 11 and 43 times longer, respectively, than those of the 1xCy3-dsDNA (median = 50 s and 8 s, respectively) (Fig. 1H and I). Furthermore, the 6xCy3-FTOB exhibited remarkable photostability under an oxygen scavenging system. During observation periods of 2000 seconds at a frame rate of 5 frames per second (fps), only 5.4% of the samples exhibited blinking behavior, and just 0.3% underwent photobleaching (Fig. S1C).

We next analyzed the 5xCy3-FTOB pair. The photobleaching step analysis was conducted on the 5xCy3-FTOB pair. The observed number of photobleaching steps closely matched the designed number of dyes, indicating that most of the samples were dimers (Fig. S1D and E). To confirm one-to-one coupling of the 5xCy3-FTOB-A and 5xCy3-FTOB-a, we prepared the FTOB-A containing five Cy5 dyes (5xCy5-FTOB-A) and mixed it with 5xCy3-FTOB-a (Fig. S1F). We observed 91% co-localization of the 5xCy5-FTOB-A and 5xCy3-FTOB-a, indicating that they were primarily heterodimers (Fig. S1G and H). Regarding the fluorescent properties of the 5xCy3-FTOB pair, the fluorescent intensity (median = 7.2 x 10^4^ a.u.) was approximately twice that of the 6xCy3-FTOB (Fig. 1F, G, and J). However, its lifetime (median = 787 s) and first blinking time (median = 471 s) were only slightly longer than those of the 6xCy3-FTOB (Fig. 1H and I). These results suggest that interaction and/or resonance energy transfer between the FTOBs did not occur due to the sufficient distance between them (approximately 10 nm) ^35^. We concluded that the 5xCy3-FTOB pair could serve as an effective probe for analyzing the collective movements of molecular motors, with minimal blinking and photobleaching.

### Versatile ALFA-tag and its nanobody effectively bind FTOB to kinesin

For the assembly of the FTOB and kinesin, we utilized the ALFA system, which includes the rationally designed ALFA-tag and a set of highly versatile nanobodies (NbALFA) that bind to the ALFA-tag with extraordinary specificity ^36^. The high-affinity NbALFA binds tightly to ALFA-tagged proteins (K_d_ = 26 pM) with a very slow off-rate (k_off_ = 9.4 × 10^−6^ s) ^36^. To bind one FTOB to one kinesin, we purified a truncated KIF5C heterodimer, with one subunit containing an NbALFA and the other a SNAP-tag (Fig. 2A and S2). The 6xCy3-FTOB was prepared using the ALFA-peptide-conjugated staple. After incubating the 6xCy3-FTOB with KIF5C, a clear band shift was observed in agarose gel electrophoresis, indicating stable binding between the FTOB and kinesin (Fig. 2B).

**Figure 2.**
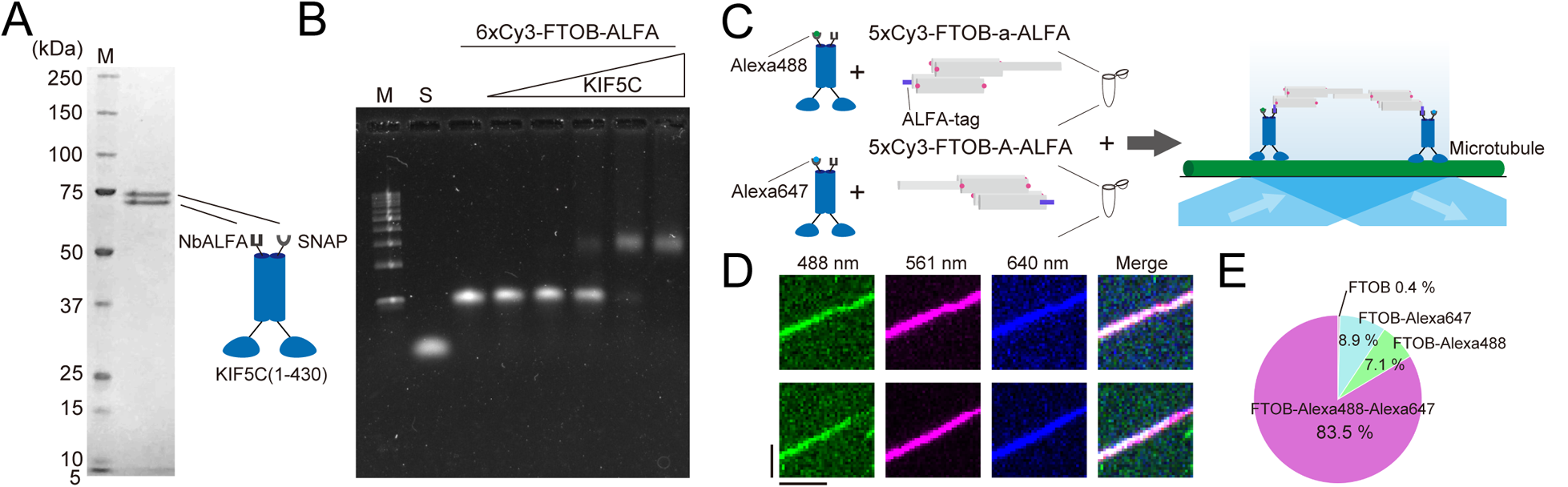
Assembly of FTOB and kinesin using the ALFA/nbALFA system. (A) SDS-PAGE analysis of a heterodimer composed of truncated KIF5C with SNAP-tag and KIF5C with nbALFA. M represents the marker lane. Numbers on the left indicate molecular weight (kDa). (B) Electrophoretic mobility shift assay. The 6xCy3-FTOB with the ALFA-tag conjugated staple was used. Lane 1: ladder marker, lane 2: scaffold DNA, lane 3: 6xCy3-FTOB, lanes 4-8: 6xCy3-FTOB and KIF5C. The concentration of KIF5C doubled at each step from lanes 4 through 8. (C) Schematic illustration of the method for binding two different motors. First, Alexa488-labeled KIF5C and Alexa647-labeled KIF5C were incubated with the 5xCy3-FTOB-a and 5xCy3-FTOB-A, respectively. Second, the two incubation solutions were mixed and incubated together. Finally, the mixture was flowed into a glass chamber and observed by TIRF microscopy. (D) Kymographs showing two representative traces of the 5xCy3-FTOB pair and KIF5C complex. Images of KIF5C-Alexa488, the 5xCy3-FTOB pair, and KIF5C-Alexa647 were alternately captured in the 488, 561, and 640 channels, respectively, and then merged. Vertical and horizontal bars represent 2 μm and 60 s, respectively. (E) Pie charts showing the relative population of each co-movement. Only 0.4% of FTOB showed no co-movement, likely due to FRET between different dyes or unlabeled motors. n = 187 (FTOB-Alexa488-Alexa647), 16 (FTOB-Alexa488), 20 (FTOB-Alexa647), and 1 (FTOB).

We next analyzed the motility of KIF5C labeled with the FTOB. To confirm that FTOB-labeled KIF5C behaves similarly to KIF5C labeled with a single dye (Alexa546), we compared their velocity and run length. The results showed nearly identical values, indicating that the FTOB does not disrupt protein function (Fig. S3). Bridging two motors with the 5xCy3-FTOB pair increased the run length but did not affect the velocity, consistent with previous studies ^30–32^ (Fig. S3).

To create a transport system driven by two different motors, we tested whether two different kinesins could be linked using the ALFA system and FTOBs (Fig. 2C). SNAP-tagged KIF5C was labeled with Alexa488 or Alexa640 and incubated with 5xCy3-FTOB-a or 5xCy3-FTOB-A, respectively. After subsequent incubation of KIF5C-Alexa488 with 5xCy3-FTOB-a and KIF5C-Alexa640 with 5xCy3-FTOB-A, the movements of the motor pairs on the microtubule were observed (Fig. 2D). The results showed that 83.5% of the 5xCy3-FTOB pairs co-moved with Alexa488 and Alexa647 (Fig. 2E), indicating that the ALFA system and FTOBs can effectively bridge two different motors.

### FTOB traces disease-associated mutant kinesins

Since we are interested in the inhibition of collective transport caused by disease-associated mutant kinesin, we introduced the E237K mutation into the *kif5c* cDNA. The E237K mutation has been identified in patients with symptoms such as epilepsy, microcephaly, and cortical malformation ^37,38^. Because these patients are heterozygous ^37,38^, it is expected that three types of dimeric KIF5C—WT/WT homodimer, E237K/E237K homodimer, and WT/E237K heterodimer—exist in the same cell. Although another disease-associated mutant, E237V, completely disrupts the ATPase activity of KIF5C ^39^, the biochemical properties of the E237K mutant have not yet been studied. In this study, we purified the E237K/E237K homodimer and WT/E237K heterodimer with NbALFA (Fig. 3A) and examined their single-molecule motilities with the FTOB. The FTOB allowed us to observe E237K/E237K homodimers and WT/E237K heterodimers on microtubules for 60 seconds or more without blinking and photobleaching (Fig. 3B and S4A). Interestingly, while the E237K/E237K homodimer was immotile and remained fixed at one point on the microtubule, the WT/E237K heterodimer moved processively (Fig. 3B and S4A).

**Figure 3.**
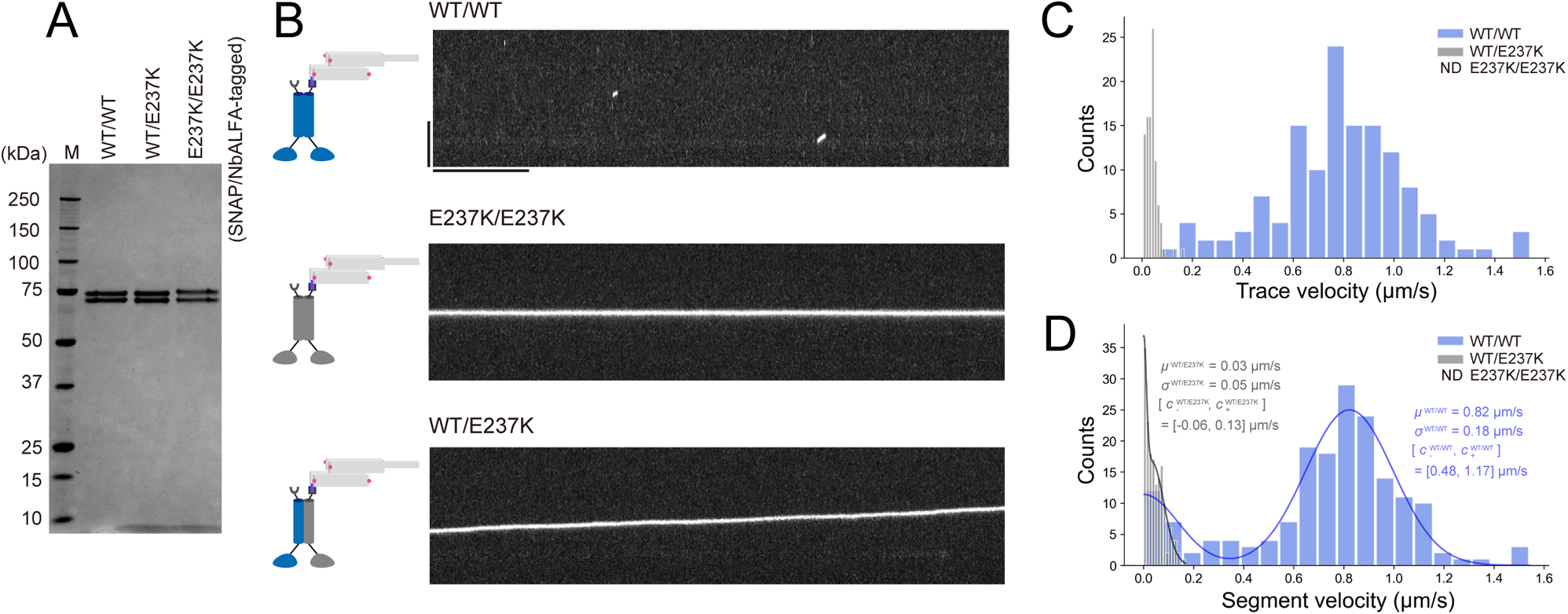
Analysis of disease-associated mutants using FTOB. (A) SDS-PAGE analysis of truncated KIF5C and its mutants. M represents the marker lane. Numbers on the left indicate molecular weight (kDa). (B) Representative kymographs showing the motility of KIF5C WT/WT homodimer, E237K/E237K homodimer, and WT/E237K heterodimer labeled by the 5xCy3-FTOB-A or 5xCy3-FTOB-a. Vertical and horizontal bars represent 5 μm and 10 s, respectively. (C) Histogram of trace velocities of the WT/WT homodimer and WT/E237K heterodimer. Note that no processive movement was detected for the E237K/E237K homodimer. n = 134 (WT/WT homodimer) and 96 (WT/E237K heterodimer). The trace velocity of the WT/WT homodimer is the same as that of KIF5C labeled with 5xCy3-FTOB-A in Fig. S3B. (D) Histogram of segment velocities of the WT/WT homodimer and WT/E237K heterodimer. The mean value (µ) and standard deviation (er) were obtained by fitting the data to the sum of two Gaussian functions, one of which has a mean of zero.

We analyzed the trace velocity by calculating it from the coordinates of the starting and ending points of the run on the microtubule. We found that the trace velocity of the WT/E237K heterodimer was significantly reduced compared to that of the WT/WT homodimer (Fig. 3C). Owing to the minimal blinking properties of the FTOB, we were able to effectively segment each trace into segments of constant velocity (Fig. 3D). While the segment and trace velocity distributions of the WT/WT homodimer were nearly identical, those of the WT/E237K heterodimer differed significantly (Fig. 3C and D). The WT/E237K heterodimer frequently paused during its run (Fig. 3D), suggesting that the E237K head often acts as an anchor, completely stalling the wild-type head on the microtubule. The FTOB proved effective in dissecting the movements of the disease-associated mutant.

### FTOB pair dissects the effect of mutant kinesin on collective transport

In the patient’s cells, the WT/WT homodimer, E237K/E237K homodimer, and WT/E237K heterodimer are expected to bind to the same cargo. Do rigor mutants like the E237K/E237K homodimer immobilize the cargo in the cell body, or do they allow wild-type motors to transport it to the axon? It is crucial to examine how the E237K/E237K homodimer and WT/E237K heterodimer disrupt the movement of the wild-type motor on cargo in vitro. The pair of 5xCy3-FTOBs was used to couple the WT/WT homodimer with the E237K/E237K homodimer or the WT/E237K heterodimer, and the movements of these motor pairs were observed (Fig. 4A, B, and S4A). As a previous study has shown that an excessive amount of rigor mutant motors can become obstacles to processive motors ^40^, we diluted the motors until the density of mutant motors on microtubules was low enough (0.30 ± 0.14 and 0.25 ± 0.09 molecules/μm for the E237K/E237K homodimer and WT/E237K heterodimer, respectively, mean ± SD) to prevent non-labeled mutant motors from interfering with the motor pairs ^40^ (Fig. S4B). Interestingly, 78% of the E237K/E237K-WT/WT pairs showed processive movement (Fig. 4A, C, and S4A). The trace velocity of the WT/E237K-WT/WT pairs was also significantly faster than that of the WT/E237K heterodimers (Fig. 3C, 4D, and S4C). However, the trace velocities of each pair were still much slower than those of the WT/WT homodimers (Fig. 3C, 4C, and D), indicating that the E237K/E237K homodimer and WT/E237K heterodimer significantly hindered the movement of the WT/WT homodimer on the cargo.

**Figure 4.**
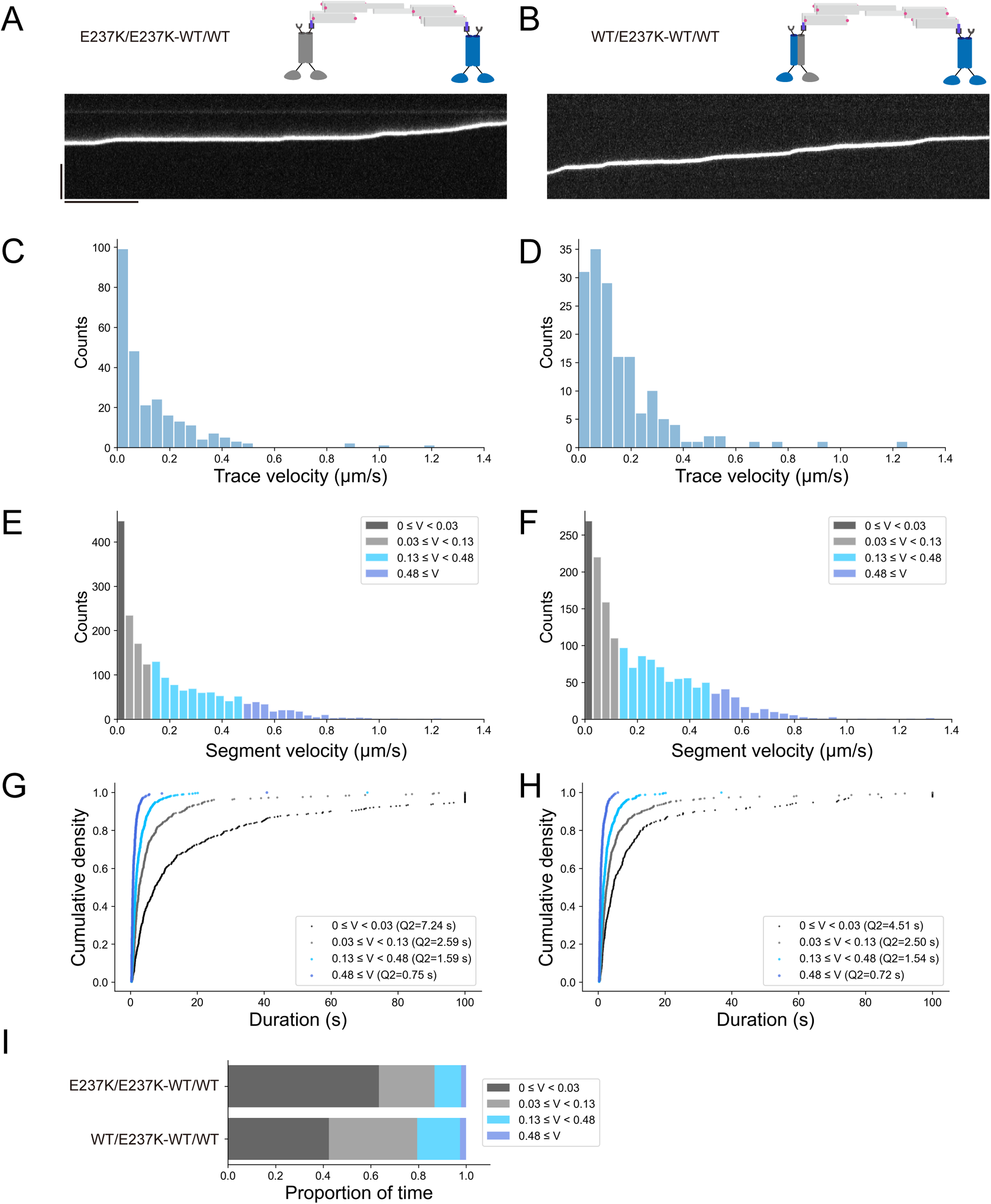
Analysis of the effect of disease-associated mutants on collective transport using FTOBs. (A and B) Representative kymographs showing the motility of (A) E237K/E237K-WT/WT pair and (B) WT/E237K-WT/WT pair. The two motors were linked via the 5xCy3-FTOB pair. Vertical and horizontal bars represent 5 μm and 10 s, respectively. (C and D) Histograms of trace velocities of (C) E237K/E237K-WT/WT pair and (D) WT/E237K-WT/WT pair. n = 257 (E237K/E237K-WT/WT pair) and 164 (WT/E237K-WT/WT pair). (E and F) Histograms of segment velocities of (E) WT/E237K-WT/WT pair and (F) WT/E237K-WT/WT pair. The segment velocities were grouped based on the fitting results of the WT/WT homodimer and WT/E237K heterodimer (Fig. 3D). (G and H) Cumulative distribution functions of trace durations in each speed section for (G) the WT/WT-E237K/E237K pair and (H) the WT/WT-WT/E237K pair. Median values (Q2) are reported. Trace durations are slightly underestimated due to limited recording time and microtubule length (e.g., when recording stops while motors are still moving or when motors reach the end of a microtubule). (I) Proportion of time spent in each speed section.

The FTOBs enabled us to clearly record velocity changes during their runs for 60 seconds or more without blinking and photobleaching (Fig. 4A, B, and S4A). The segment velocities of the E237K/E237K-WT/WT pairs and WT/E237K-WT/WT pairs were categorized based on the mean segment velocity (µ) and standard deviation (er) of the WT/WT homodimer and WT/E237K heterodimer (Fig. 3D, 4E and F). The velocity ranges were defined as stationary (0<*v* < 0.03 µm/s), slow (0.03 < *v* < 0.13 µm/s), medium (0.13 <*v* < 0.48 µm/s), and fast (v > 0.48 µm/s), corresponding to 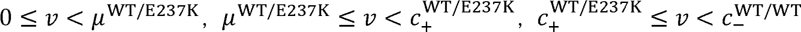, and 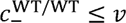, respectively. Here, 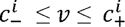 (*i* = WT/WT or WT/E237K) denotes the range within which 95% of the data for motor i is predicted to fall 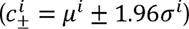. The duration times within these four velocity ranges during the runs were measured (Fig. 4G, H), and the proportions of time spent in each range were calculated (Fig. 4I). During the fast speed range, which accounted for about 2% of the time, both pairs could move at the same speed as the WT/WT homodimer, suggesting that the mutant motor temporarily detaches from the microtubule, allowing the wild-type motor to move smoothly (Fig. 4I). However, this period was very short-lived (Fig. 4G, H). One might think this duration was due to the wild-type motor not being linked to the mutant motor, but this is unlikely since the proportion of fast-speed duration calculated directly from the trace velocities was negligible (about 0.3%) (Fig. S4D). For 34% of the time (slow and medium speed ranges), the E237K/E237K-WT/WT pairs moved unidirectionally. For 18% of the time (medium speed range), the velocity of the WT/E237K-WT/WT pairs was significantly faster than that of the WT/E237K heterodimer. During these periods, the wild-type motor likely dragged its mutant partner along the microtubule. For the remaining time, their speeds were almost consistent with those of the mutant motors, implying that the wild-type motor was completely suppressed by the mutant. The detailed inhibitory effects of the mutant motors were successfully characterized by our system using the FTOBs.

## DISCUSSION

### Advantage of FTOB

In this study, we developed a highly photostable DNA origami block that can bind to a protein of interest. The previously developed FluoroCube, based on the single-stranded tile method ^35^, requires strict concentration control of each single-stranded DNA to avoid undesired misfolded and/or defective structures. In our DNA origami-based approach, each functionalized staple DNA is designed to hybridize only with the scaffold strand, ensuring efficient formation of the target structure, as reflected by the presence of a single main band (except for the one corresponding to free staple DNAs) in AGE. Owing to its minimal components and simple purification process, the FTOB is easier to prepare compared to typical DNA origami structures that require many components ^31^. Another advantage of our FTOB is its ability to hybridize with a partner FTOB, enabling the coupling of two different molecular motors and the observation of their collective movements without frequent blinking and photobleaching. As demonstrated in the observation of wild-type and mutant motor pairs, our system is particularly effective for capturing rare, short-lived events that require long recording times at high FPS, where conventional single dyes often undergo blinking and photobleaching.

### ALFA-tag/NbALFA system for assembling DNA nanostructures and proteins

We introduced the ALFA-tag/NbALFA system for assembling DNA nanostructures and proteins. Since NbALFA is a small and stable protein, it can be expressed as a fusion protein in bacteria. Typically, antigen-antibody reactions are not suitable for assembling proteins with DNA nanostructures. Most antigens and antibodies are too large to be modified for DNA, and even if they can be modified, the high-temperature annealing process required for folding DNA origami likely deactivates them. The ALFA-tag, a short α-helix (13 amino acids) ^36^, retains its strong and stable binding to NbALFA even after annealing at 85 °C, as shown by a band shift in AGE.

Covalent SNAP-tag protein technology has generally been used for assembling proteins and DNA nanostructures. However, major challenges with the SNAP-tag include slow assembly speed (k_on_ = 6 × 10^3^ M^−1^s^−1^) and low integration yield between negatively charged DNA nanostructures and kinesin with affinity tags, likely due to charge repulsion ^41^. Given the very fast on-rate (k_on_ = 3.6 × 10^5^ M^−1^s^−1^) and slow off-rate (k_off_ = 9.4 × 10^−6^ s^−1^) between the ALFA-tag and NbALFA ^36^, it may be possible to attach proteins of interest to larger DNA origami structures quickly and stably using the ALFA-tag/NbALFA system.

### Inhibition of transport caused by KIF5C(E237K) mutation

Despite the differences in motility between the E237K/E237K homodimer and WT/E237K heterodimer, both similarly inhibited the wild-type motor during collective transport. The force-dependence of motor detachment is a crucial factor in determining speed in collective transport ^42–44^. Most kinesins detach more easily from the microtubule in the ADP state than in the ATP state under load ^45^. In the case of KIF5B(E236A), the neck linker is stabilized in an ATP-like, docked conformation, allowing the motor to bind strongly to the microtubule at all times ^46^. The same may be true for KIF5C(E237K). Indeed, most of the time, both the E237K/E237K homodimer and WT/E237K heterodimer overpowered the wild-type motor (Fig. 4). However, our system clearly detected that the wild-type motor occasionally managed to handle the load from the mutant motor, likely causing the mutant motor to either detach or be dragged during collective transport.

In the patient’s cells, if both wild-type and mutant monomers are expressed equally, the ratio of wild-type/wild-type homodimers, wild-type/mutant heterodimers, and mutant/mutant homodimers on cargo vesicles is expected to be 1:2:1. Therefore, the inhibition of wild-type motors by the mutants may be even more severe than observed in this study. This motor ratio can be replicated by further developing FTOBs for trimerization and tetramerization. The advantage of our approach is that it does not require different tags to combine various types of motors, making it easy to test different motor type ratios on cargo. This system will provide further insights into intracellular transport and its deficiencies, particularly the coordination between different types of motors and their mutants.

## METHODS

### Preparation of FTOBs

Fluorescence-labeled and unmodified DNA strands were purchased from Eurofins Genomics Tokyo (Tokyo, Japan) or Integrated DNA technologies (Coralville, IA). Peptide-modified oligonucleotide was purchased from bioSYNTHESIS (Lewisville, TX). FTOB structures were designed using the caDNAno software ^47^ for strand routing. Assembly of the FTOB structures was accomplished by mixing 84 nt of scaffold DNA with staple strands to a final concentration of 10 μM in 25 μL of the folding buffer containing 5 mM Tris-HCl (pH8.0), 1 mM EDTA, and 15 mM MgCl_2_ (Table S1 and 2). The mixture was incubated at 85 °C for 5 min, then annealed by gradually lowering the temperature from 80 °C to 65 °C at a rate of −1 °C per 5 min, followed by a reduction from 65 °C to 25 °C at a rate of −1 °C per 20 min, and finally held at 4 °C. The assembled FTOB structures were analyzed by agarose gel electrophoresis and then purified by gel excision followed by centrifugation in Freeze’N Squeeze columns (Bio-Rad Laboratories, Inc. Hercules, CA).

### Agarose gel electrophoresis

The samples were loaded for electrophoresis on a 3.0% agarose gel containing 5 mM MgCl_2_ in a 0.5× Tris-borate-EDTA buffer and then electrophoresed in 0.5× TBE with 5 mM MgCl_2_ under 90 V at 4 °C. The gels were then imaged with iBright FL1500 Imaging System (Thermo Fisher Scientific, Waltham, MA) using SYBR Gold Nucleic Acid Gel Stain (Thermo Fisher Scientific, Waltham, MA) as the staining dye.

### Plasmid of kinesin

The pET-32a vector containing *Rattus norvegicus* KIF5C(1-430)::SNAP::His and KIF5C(1-430)::FLAG was provided by Dr. Ken’ya Furuta ^32^. In this construct, cysteine 7 in KIF5C was mutated to serine. As previously reported, this mutation did not affect its motility ^46^, so we referred to this mutant as the wild-type in this study. We replaced the FLAG tag with NbALFA linked to a Strep II tag. The mutant KIF5C was generated through PCR-based mutagenesis.

### Purification of kinesin

Reagents were purchased from Nacarai tesque (Kyoto, Japan), unless described. BL21(LOBSTR) was transformed and selected on LB agar supplemented with ampicillin at 37 °C overnight. Colonies were picked and cultured in 5 ml LB medium supplemented with ampicillin overnight. Next morning, 5 ml of the medium was transferred to 1000 ml 2.5× YT (20 g/L Tryptone, 12.5 g/L Yeast Extract, 6.5 g/L NaCl) supplemented with 10 mM phosphate buffer (pH 7.4) and 50 μg/ml ampicillin in a 2 L flask and shaken at 37 °C. Two flasks were prepared. When OD600 reached 0.6, flasks were cooled in ice-cold water for 30 min. Then, 60 mg IPTG was added to each flask. Flasks were shaken at 18°C overnight. Next day, bacteria expressing recombinant proteins were pelleted by centrifugation (3000 g, 10 min, 4 °C), resuspended in PBS and centrifuged again (3000 g, 10 min, 4°C). Pellets were resuspended in protein buffer (50 mM Hepes, pH 7.4, 150 mM KCH_3_COO, 2 mM MgSO_4_, 1 mM EGTA, 10% glycerol) supplemented with Phenylmethylsulfonyl fluoride (PMSF). Bacteria were lysed by sonication. Lysate was obtained by centrifugation (75,000 g, 20 min, 4 °C). Lysate was loaded on Streptactin-XT resin (IBA Lifesciences, Göttingen, Germany) (bead volume: 2 ml). The resin was washed with 40 ml wash buffer (50 mM Hepes, pH 7.4, 500 mM KCH_3_COO, 2 mM MgSO_4_, 1 mM EGTA, 10% glycerol). Protein was eluted with 30 ml Strep elution buffer (50 mM Hepes, pH 7.4, 150 mM KCH_3_COO, 2 mM MgSO_4_, 1 mM EGTA, 10% glycerol, 50 mM biotin). Eluted solution was then loaded on TALON resin (Takara Bio Inc., Kusatsu, Japan) (bead volume: 4 ml). The resin was washed with 40 ml wash buffer and eluted with His-tag elution buffer (50 mM Hepes, pH 7.4, 150 mM KCH_3_COO, 2 mM MgSO_4_, 10% glycerol, 500 mM imidazole). Eluted solution was concentrated using an Amicon Ultra 15 and then separated on an NGC chromatography system (Bio-Rad) equipped with a Superdex 200 Increase 10/300 GL column (Cytiva). Peak fractions were collected, and some were labeled with SNAP-Surface Alexa Fluor 488, 546, or 647 (NEB, S9129S, S9132S, S9136S) according to the manufacturer’s instructions. The samples were then aliquoted and snap-frozen in liquid nitrogen.

### Preparation of microtubules

Tubulin was purified from porcine brain as described ^48^. Tubulin was labeled with Biotin-PEG2-NHS ester (Tokyo Chemical Industry, Tokyo, Japan) and AZDye647 NHS ester (Fluoroprobes, Scottsdale, AZ, USA) as described ^49^. To polymerize Taxol-stabilized microtubules labeled with biotin and AZDye647, 30 μM unlabeled tubulin, 1.5 μM biotin-labeled tubulin, and 1.5 μM AZDye647-labeled tubulin were mixed in BRB80 buffer (80 mM PIPES, 1mM MgCl_2_, and 1mM EGTA) supplemented with 1 mM GTP and incubated for 15 min at 37 °C. Then, an equal amount of BRB80 supplemented with 40 μM taxol was added and further incubated for more than 15 min. The solution was loaded on BRB80 supplemented with 300 mM sucrose and 20 μM taxol and ultracentrifuged at 100,000×g for 5 min at 30 °C. The pellet was resuspended in BRB80 supplemented with 20 μM taxol. In the measurement of co-movement of the 5xCy3-FTOB pair, KIF5C-Alexa488, and KIF5C-Alexa647, the microtubules were not fluorescently labeled.

### Preparation of flow cells with dsDNA and FTOBs

Glass chambers were prepared by acid washing as previously described ^50^. The chambers were then coated with PLL-PEG-biotin (SuSoS, Duubendorf, Switzerland) followed by streptavidin. Unbound streptavidin was washed away using FTOB assay buffer (the folding buffer with 0.5 mg/ml kappa-casein and 0.5% Pluronic F127). To form dsDNA, 5 µM Cy3-labeled ssDNA (ssDNA-Cy3) was annealed to 5 µM biotin-labeled ssDNA (FTOB_0_Bio) at 25 °C for 1 hour, following an incubation at 85 °C for 5 min (Table S1 and 2). For the 5xCy3-FTOB pair, the 5xCy3-FTOB-A and 5xCy3-FTOB-a (final concentration, 10 nM each) were incubated in the folding buffer at 25 °C for approximately 60 minutes. Either dsDNA, FTOB, or FTOB pair was then diluted to 10 pM in FTOB assay buffer and flowed into streptavidin-coated flow chambers. Unbound dsDNA, FTOB, or FTOB pair was washed away using FTOB assay buffer. In measurement under the oxygen scavenger system (Fig. S1C), 2 mM protocatechuic acid (PCA), 125 μM protocatechuate-3,4-dioxygenase (PCD), and 2 mM Trolox were included in the FTOB buffer. In the photobleaching step analysis, 100 μM PCA, 6.25 μM PCD, and 100 μM Trolox were included in the FTOB buffer. In this analysis, a pair of 4xCy3-FTOB-A and 4xCy3-FTOB-a was used instead of the 5xCy3-FTOB pair to more accurately detect the number of photobleaching steps.

### Preparation of flow cells with kinesin

Glass chambers coated with PLL-PEG biotin and streptavidin were prepared as described in the section ‘*Preparation of flow cells with dsDNA and FTOBs’*. Unbound streptavidin was washed away with BRB80. Polymerized microtubules were then introduced into the streptavidin-coated flow chambers and allowed to adhere for 5–10 minutes. Unbound microtubules were removed using SRP90 assay buffer [90 mM Hepes pH 7.4, 50 mM KCH_3_COO, 2 mM Mg(CH_3_COO)_2_, 5 mM MgCl_2_, 1 mM EGTA, 10% glycerol, 0.1 mg/ml biotin–BSA, 0.2 mg/ml kappa-casein, 0.5% Pluronic F127, ATP (2 mM or 5 µM), and an oxygen scavenging system composed of 2 mM PCA/125 µM PCD/2 mM Trolox]. Two different or identical motors (motor A and motor B) were linked as follows: 5xCy3-FTOB-A (at a final concentration of 30 nM) and motor A (at a final concentration of 300 nM) were incubated in folding buffer at 25 °C for approximately 45 minutes. The same incubation was performed for 5xCy3-FTOB-a and motor B. The two incubation solutions were then mixed and incubated at 25 °C for approximately 40 minutes. The FTOB-motor complex was diluted to 300 pM in SRP90 assay buffer unless otherwise specified. The solution was then introduced into the glass chamber. In tracking the co-movement of FTOBs, KIF5C-Alexa488, and KIF5C, the assay was conducted in the presence of 5 µM ATP. Other assays were performed in the presence of 2 mM ATP. Because the concentrations of the FTOB-motor complex mentioned above were not sufficient to measure a significant number of runs for the single wild-type motor or identical wild-type pairs in the presence of 2 mM ATP (where they were short-lived on microtubules), the concentration was increased tenfold. Since an excessive amount of free rigor mutants can obstruct processive motors ^40^, the mutant motors were labeled with Alexa488, and their density on the microtubules was examined in the 488 channel (Fig. S4B).

### Single-molecule TIRF data collection and analysis of dsDNA and FTOBs

An ECLIPSE Ti2-E microscope equipped with a CFI Apochromat TIRF 100XC Oil objective lens, an Andor iXion life 897 camera, and a Ti2-LAPP illumination system (Nikon, Tokyo, Japan) was used to observe single-molecule motility. NIS-Elements AR software ver. 5.2 (Nikon) was used to control the system. The TIRF data of the surface-immobilized dsDNA, FTOB, or FTOB pair was acquired under continuous laser illumination. 7500 frames (corresponding to approximately 1500 s) for the dsDNA and 10000 frames (corresponding to approximately 2000 s) for the FTOB and FTOB pair were taken at a frame rate of 5 fps in the 561 channel unless otherwise specified. In observation of the colocalization of the FTOBs, images of 5xCy3-FTOB-a and 5xCy5-FTOB-A were alternately taken at a frame rate of 0.31 fps in the 561 and 640 channels, respectively. All experiments were performed at room temperature (25 °C) at least three times independently. The data was analyzed in ImageJ. The Subtract Background tool (10-pixel radius, sliding paraboloid) was used to subtract background fluorescence from each frame in the stack. Single molecules were located and traced using the Spot Intensity Analysis plugin in ImageJ with the following settings: time interval of 0.2 s, electrons per analog-to-digital unit (ADU) of 1.84, spot radius of 6, noise tolerance of 500 (except for the dsDNA for which a noise tolerance of 300 was used), and a median background estimation. The number of frames to check is 10 frames. To quantify the fluorescence intensity of each molecule, the average fluorescence intensity over each set of 20 frames was calculated, and the highest average value was taken as the representative fluorescence intensity. A probe was considered photobleached if it remained in a dark state for more than 50 seconds; all other events were classified as blinking. For the photobleaching step analysis, each photobleaching point was identified by visual inspection.

### Single-molecule TIRF data collection and analysis of kinesin

The microscope and setup were the same as described in ‘*Single-molecule TIRF data collection and analysis of dsDNA and FTOBs*.’ TIRF data of kinesin on microtubules was acquired under continuous laser illumination. A total of 1000 frames (corresponding to approximately 100 seconds) were captured at a frame rate of 10 fps in the 561 channel, unless otherwise specified. In observation of the co-movement, images of KIF5C with Alexa488, the 5xCy3-FTOB pair, and KIF5C with Alexa640 were alternately captured at a frame rate of 0.3 fps in the 488, 561, and 640 channels, respectively. All experiments were performed at room temperature (25 °C) at least three times independently. The data was analyzed using kymographs in ImageJ. Motors with run lengths of 300 nm (corresponding to 3 pixels) or greater were counted as processive and included in the analysis, while others were excluded. Velocity segmentation of full traces was done manually, requiring a velocity change of more than 30 nm/s, with each segment lasting at least three frames (approximately 300 ms) to be considered. The mean segment velocity (µ) and standard deviation (er) of the WT/WT homodimer and WT/E237K heterodimer were determined by fitting the segment velocities to a sum of two Gaussian functions, N(µ, er^2^) and N_0_(0, er^2^). The proportion of time spent in each speed section was calculated by dividing the total time in each section by the overall time.

### Statistical analyses and graph preparation

Statistical analyses were performed using Graph Pad Prism version 9. Statistical methods are described in the figure legends. Graphs were prepared using python or Graph Pad Prism version 9, and aligned by Adobe Illustrator 2021.

## ACKNOWLEDGMENTS

We thank Dr. K. Furuta for providing the original plasmid of KIF5C kinesin and Dr. I. Kawamata for assisting the sequence design of connector DNAs for FTOBs. We would also like to thank the members of the Niwa lab and Tsumoto lab for their valuable discussions. T.K. was supported by JSPS KAKENHI (grant no. JP23KJ0168). S.N. was supported by JSPS KAKENHI (grant no. JP23H02472). Y.S. was supported by JSPS KAKENHI (grant no. JP23K24937, JP23H04416, and JP23K17860).

## AUTHOR CONTRIBUTIONS

Conceptualization, Y.S. and S.N; Methodology, Y.S. and S.N.; Investigation, T.K., R.S., Y.S., and S.N.; Writing – Original Draft, T.K.; Writing – Review & Editing, Y.S. and S.N.; Funding Acquisition, T.K., Y.S., and S.N.; Resources, T.K., R.S., Y.S., and S.N.; Supervision, Y.S., and S.N.

## DECLARATION OF GENERATIVE AI AND AI-ASSISTED TECHNOLOGIES

During the preparation of this work the authors used ChatGPT4o in order to check English grammar and improve English writing. After using this tool, the authors reviewed and edited the content as needed and take full responsibility for the content of the publication.

## SUPPLEMENTAL INFORMATION

Supplemental Information: Figures S1–S4

