## Supplemental Information for "Modular photostable fluorescent DNA blocks dissect the effects of pathogenic mutant kinesin on collective transport"

### Supporting Tables

Table S1: Sequences and corresponding modifications of all oligonucleotides used in this study.

| Name | Sequence and modification |
|---|---|
| FTOB_Scaff | CTCAGGATTATCAACCGGGGTACATATGATTGACATGCTAGTTTTACGATTACCGTTCATCG ATTCTCTTGTTTGCTCCAGACT |
| FTOB_0_Bio | 5Biosg/GTAAAACTAGCATGCAAACAA |
| FTOB_0_ALFA | 5ALFA peptide (SRLEEELRRRLTE)/GTAAAACTAGCATGCAAACAA |
| FTOB_1_Cy3 | 5Cy3/TATGTACCATGAACGGTAATC/3Cy3Sp |
| FTOB_1_Cy5 | 5Cy5/TATGTACCATGAACGGTAATC/3Cy5Sp |
| FTOB_2_Cy3 | 5Cy3/GAGAATCGCCGGTTGATAATC/3Cy3Sp |
| FTOB_2_A | GAGAATCGCCGGTTGATAATCTTGCCCAAATAAGCCCAATAAAGTCCACAGTCCC |
| FTOB_2_a | GAGAATCGCCGGTTGATAATCTTGGGACTGTGGACTTTATTGGGCTTATTTGGGC |
| FTOB_2_Cy3_A | 5Cy3/GAGAATCGCCGGTTGATAATCTTGCCCAAATAAGCCCAATAAAGTCCACAGTCCC |
| FTOB_2_Cy3_a | 5Cy3/GAGAATCGCCGGTTGATAATCTTGGGACTGTGGACTTTATTGGGCTTATTTGGGC |
| FTOB_2_Cy5_A | 5Cy5/GAGAATCGCCGGTTGATAATCTTGCCCAAATAAGCCCAATAAAGTCCACAGTCCC |
| FTOB_3_Cy3 | 5Cy3/CTGAGAGTCTGGAGTCAATCA/3Cy3Sp |
| FTOB_3_Cy5 | 5Cy5/CTGAGAGTCTGGAGTCAATCA/3Cy5Sp |
| ssDNA_Cy3 | 5Cy3/TTGTTTGCATGCTAGTTTTAC |

Table S2: Combinations of oligonucleotide strands used to assemble all DNA constructs.

| Construct | Oligonucleotide strands |
|---|---|
| 6xCy3-FTOB-Biotin | FTOB_Scaff, FTOB_0_Bio, FTOB_1_Cy3, FTOB_2_Cy3, FTOB_3_Cy3 |
| 6xCy3-FTOB-ALFA | FTOB_Scaff, FTOB_0_ALFA, FTOB_1_Cy3, FTOB_2_Cy3, FTOB_3_Cy3 |
| 5xCy3-FTOB-A-Biotin | FTOB_Scaff, FTOB_0_Bio, FTOB_1_Cy3, FTOB_2_Cy3_A, FTOB_3_Cy3 |
| 5xCy3-FTOB-a-Biotin | FTOB_Scaff, FTOB_0_Bio, FTOB_1_Cy3, FTOB_2_Cy3_a, FTOB_3_Cy3 |
| 5xCy3-FTOB-A-ALFA | FTOB_Scaff, FTOB_0_ALFA, FTOB_1_Cy3, FTOB_2_Cy3_A, FTOB_3_Cy3 |
| 5xCy3-FTOB-a-ALFA | FTOB_Scaff, FTOB_0_ALFA, FTOB_1_Cy3, FTOB_2_Cy3_a, FTOB_3_Cy3 |
| 4xCy3-FTOB-A-Biotin | FTOB_Scaff, FTOB_0_Bio, FTOB_1_Cy3, FTOB_2_A, FTOB_3_Cy3 |
| 4xCy3-FTOB-a-Biotin | FTOB_Scaff, FTOB_0_Bio, FTOB_1_Cy3, FTOB_2_a, FTOB_3_Cy3 |
| 5xCy5-FTOB-A-Biotin | FTOB_Scaff, FTOB_0_Bio, FTOB_1_Cy5, FTOB_2_Cy5_A, FTOB_3_Cy5 |
| 1xCy3-dsDNA-Biotin | FTOB_0_Bio, ssDNA_Cy3 |

### Supporting Figures

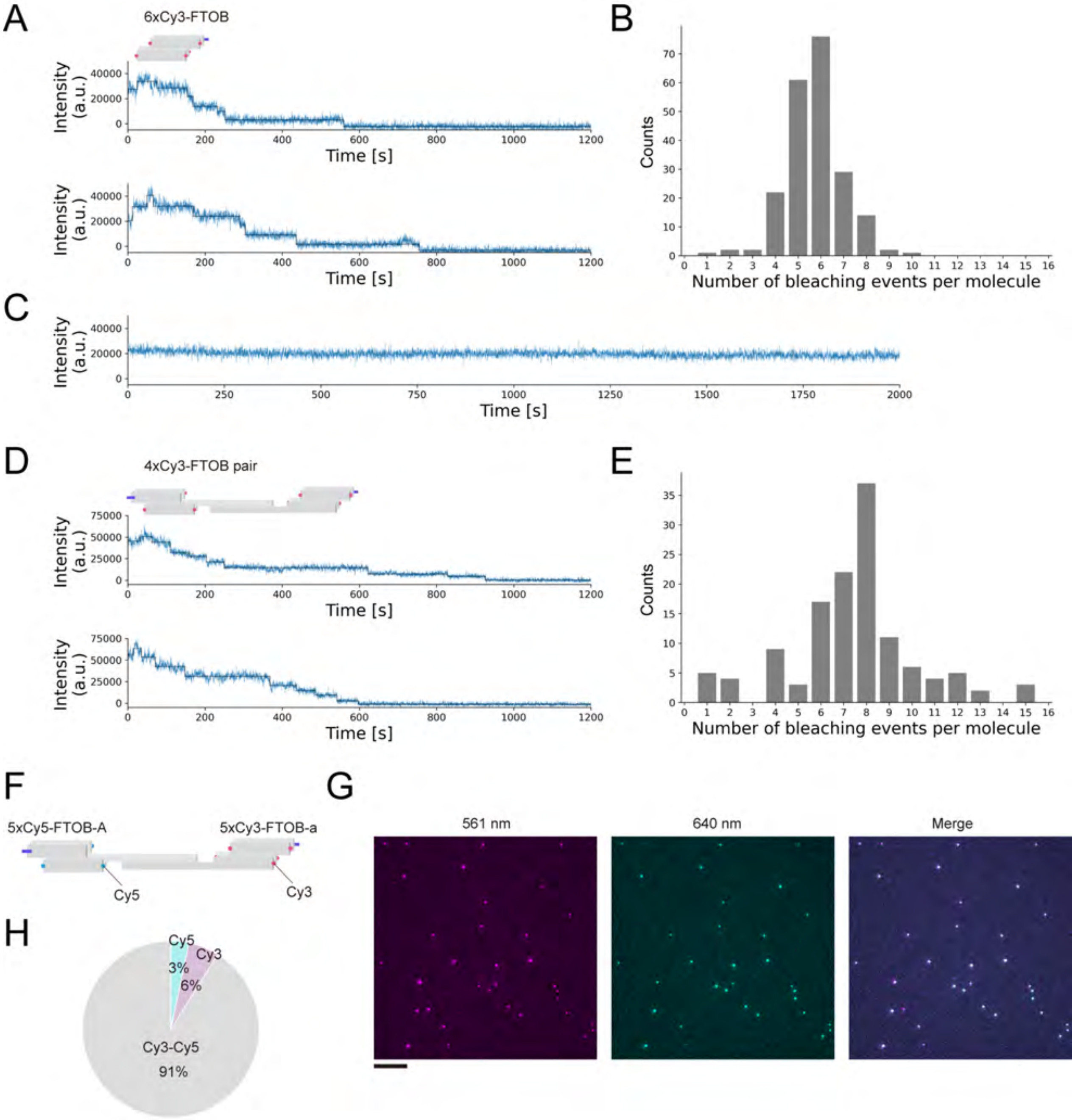


Figure S1: Further analysis of the photophysical properties of the FTOBs. (A) Example intensity traces of the 6xCy3-FTOB in the presence of 100 $\mu$M PCA, 6.25 $\mu$M PCD, and 100 $\mu$M Trolox with detected photobleaching steps. (B) Histogram of the number of photobleaching steps from the 6xCy3-FTOB. n=210. (C) Example intensity trace of the 6xCy3-FTOB in the presence of 2 mM PCA, 125 $\mu$M PCD, and 2 mM Trolox. (D) Example intensity traces of a pair of 4xCy3-FTOB-A and 4xCy3-FTOB-a in the presence of 100 $\mu$M PCA, 6.25 $\mu$M PCD, and 100 $\mu$M Trolox with detected photobleaching steps. (E) Histogram of the number of photobleaching steps from a pair of 4xCy3-FTOB-A and 4xCy3-FTOB-a. n=128. (F) Schematic illustration of a pair of 5xCy5-FTOB-A and 5xCy3-FTOB-a. (G) Example TIRF images of 5xCy5-FTOB-A and 5xCy3-FTOB-a. Images were captured in the 561 and 640 channels and then merged. Scale bar represents 10 $\mu$m. (H) Pie charts showing the relative proportions of localization and colocalization events. n=276 (Cy3-Cy5), n=17 (Cy3), and n=10 (Cy5).

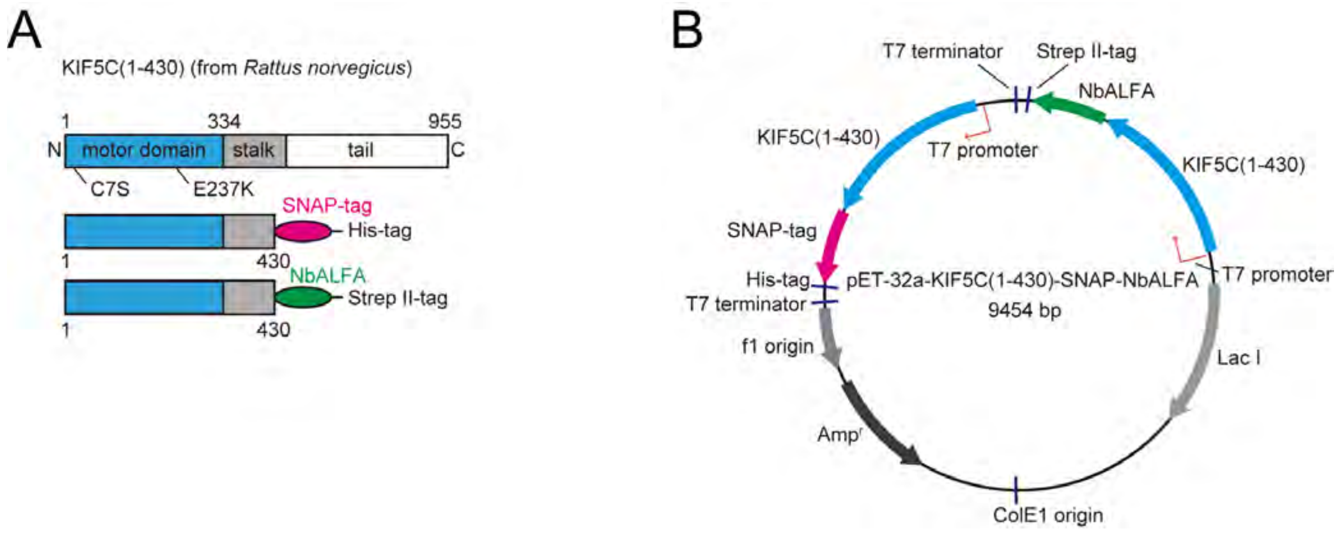


Figure S2: Construction of kinesin. (A) Schematic representations of KIF5C constructs. Full-length proteins are shown at the top, with the positions of mutations indicated. (B) Diagrams of the vectors that express the heterodimer of KIF5C.

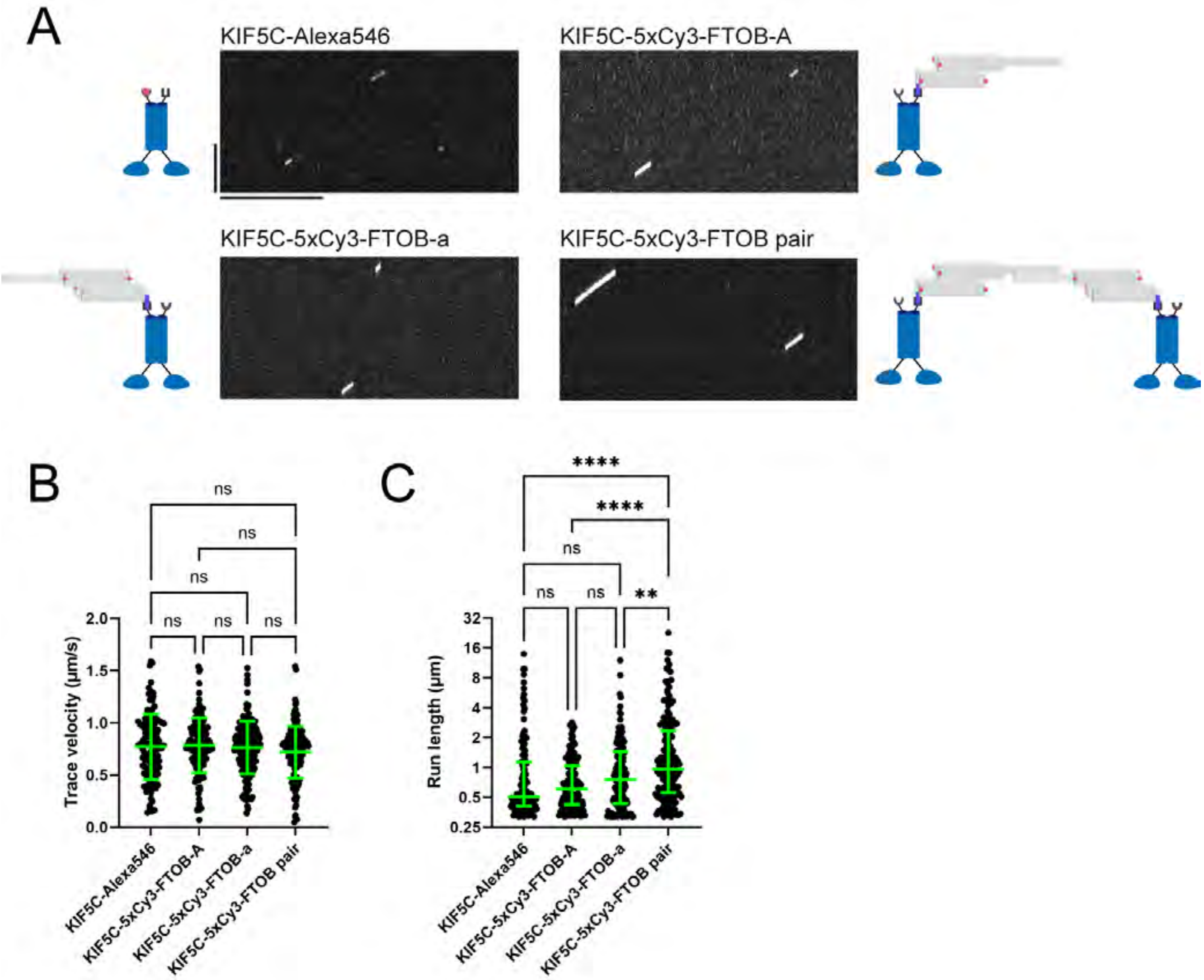


Figure S3: Comparison of motility between KIF5C labeled with a single dye (Alexa 546), 5xCy3-FTOB-A, 5xCy3-FTOB-a, and the 5xCy3-FTOB pair. (A) Representative kymographs showing the motility of KIF5C labeled with Alexa 546, 5xCy3-FTOB-A, 5xCy3-FTOB-a, or the 5xCy3-FTOB pair. Vertical and horizontal bars represent 5 $\mu$m and 10 s, respectively. (B) Dot plots showing the trace velocity of KIF5C-Alexa546, KIF5C-5xCy3-FTOB-A, KIF5C-5xCy3-FTOB-a, and KIF5C-5xCy3-FTOB pair. Green bars represent mean ± S.D. n = 119 (KIF5C-Alexa546), 134 (KIF5C-5xCy3-FTOB-A), 162 (KIF5C-5xCy3-FTOB-a), and 170 (KIF5C-5xCy3-FTOB pair). One-way ANOVA test followed by Šidák's multiple comparison test. ns, $p>0.05$ (not statistically significant). (C) Dot plots showing the run length of KIF5C-Alexa546, KIF5C-5xCy3-FTOB-A, KIF5C-5xCy3-FTOB-a, and KIF5C-5xCy3-FTOB pair. The y-axis is plotted on a logarithmic scale with base 2. Green bars represent median value and interquartile range. n = 119 (KIF5C-Alexa546), 134 (KIF5C-5xCy3-FTOB-A), 162 (KIF5C-5xCy3-FTOB-a), and 170 (KIF5C-5xCy3-FTOB pair). Kruskal–Wallis test followed by Dunn's multiple-comparison test. ns, $p>0.05$ (not statistically significant). **, $p<0.01$. ****, $p<0.0001$.

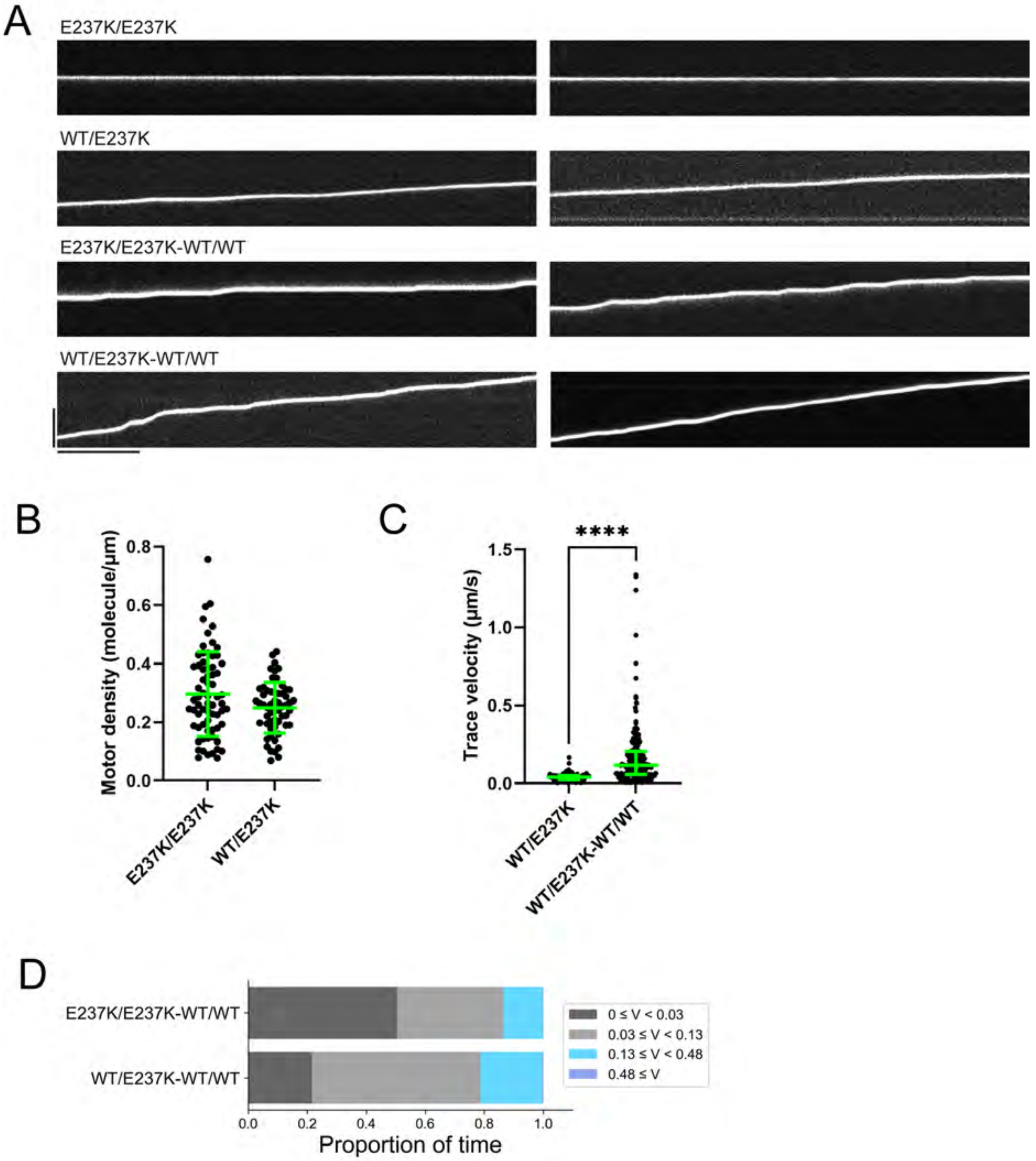


Figure S4: Further analysis of mutant motors and their pairs with wild-type motors. (A) Additional representative kymographs showing the motility of the E237K/E237K homodimer, WT/E237K heterodimer, E237K/E237K-WT/WT pair, and WT/E237K-WT/WT pair. Vertical and horizontal bars represent 5 $\mu$m and 10 s, respectively. (B) Dot plots showing the motor density of the E237K/E237K homodimer and WT/E237K heterodimer on microtubules. Green bars represent mean ± S.D. n=68 and 57 microtubules for the E237K/E237K homodimer and WT/E237K heterodimer, respectively. (C) Statistical comparison between the trace velocities of the WT/E237K heterodimer and WT/E237K-WT/WT pair. Green bars represent median value and interquartile range. The data are the same as shown in Fig. 3C and 4C. n=96 and 164 molecules for the WT/E237K heterodimer and WT/E237K-WT/WT pair, respectively. Mann-Whitney U test. ****, $p<0.0001$. (D) Proportion of time spent in each speed section calculated from the trace velocities.
